# Amygdala-predominant α-synuclein pathology exacerbates hippocampal neuron loss in Alzheimer’s disease

**DOI:** 10.1101/2024.06.21.599515

**Authors:** Klara Gawor, Sandra Tomé, Rik Vandenberghe, Philip Van Damme, Mathieu Vandenbulcke, Markus Otto, Christine A.F. von Arnim, Estifanos Ghebremedhin, Alicja Ronisz, Simona Ospitalieri, Matthew Blaschko, Dietmar R. Thal

**Author notes:** Correspondence to: Dietmar Thal, Klara Gawor ON4 Herestraat 49 - box 1032, 3000 Leuven.

## Abstract

Misfolded α-synuclein (αSyn) protein accumulates in 43-63% of individuals with symptomatic Alzheimer’s disease (AD). Two main patterns of co-morbid αSyn pathology have been identified: caudo-rostral and amygdala-predominant, with the latter being more common in AD. αSyn pathology has been shown to interact with DNA-binding protein 43 (TDP-43) and abnormally phosphorylated Tau protein (pTau). These proteins tend to accumulate in the amygdala, yet the specific role of amygdala-predominant αSyn pathology in the progression of AD and hippocampal degeneration remains unclear.

In this cross-sectional study, we analyzed 291 autopsy brains from both demented and non-demented elderly individuals neuropathologically. Neuronal density in the CA1 region of the hippocampus was assessed using hematoxylin-stained sections for all cases. We semi-quantitatively evaluated αSyn pathology severity in six brain regions and stratified the cases into the two spreading patterns. In 99 AD cases, we assessed limbic-predominant age-related TDP-43 neuropathological changes (LATE-NC), CA1 pTau density, and cerebral amyloid angiopathy (CAA). Structural equation modeling analysis was conducted based on the assessed pathological parameters in AD patients.

We identified an association between the amygdala-predominant αSyn pathology pattern and decreased neuronal density in the CA1 region. AD patients with an amygdala-predominant αSyn pattern exhibited the most severe pTDP-43 pathology among all groups, while those with the caudo-rostral pattern had the lowest severity of AD-related pathological changes including CAA type 1. Our model revealed that the relationship between αSyn pathology and CA1 neuron loss is mediated through pTau and LATE-NC.

Our results indicate that amygdala-predominant αSyn pathology, in contrast to αSyn pathology with a caudo-rostral pattern, significantly contributes to hippocampal neuron loss, potentially by accelerating TDP-43 and pTau pathologies. This finding, along with observed neuropathological differences between AD patients with these two αSyn spreading patterns, underscores the need for precise stratification of patients. The stratification should consider not only the molecular and morphological identity of co-pathologies but also the distribution pattern of the respective co-pathologies.

## Introduction

Alpha-synuclein (αSyn) is a protein abundantly present in the human brain, primarily localized within pre-synaptic terminals under normal physiological conditions^1^. The accumulation of misfolded αSyn in the gray matter is a significant contributor to age-related brain degeneration, clinically manifested as Lewy Body Disorders (LBD), which encompass Parkinson’s disease, Parkinson’s disease dementia, and Dementia with Lewy Bodies^2^. In LBD, αSyn aggregates can take the form of Lewy bodies, Lewy neurites, and astrocytic inclusions, depending on their specific locations^3^. Remarkably, similar morphological αSyn pathology has been detected in 43-63% of individuals diagnosed with Alzheimer’s disease (AD).^4–7^

In advanced cases with αSyn pathology the aggregates are observed in multiple brain regions, including the brainstem, limbic system, and neocortex.^8–11^ Although αSyn pathology in many affected individuals follows a typical pattern of distribution described by Braak,^9^ considerable heterogeneity exists in the severity across brain regions. Various attempts to identify underlying spreading patterns have been made.^12–14^ Despite some variations, a consistent trend has emerged indicating that in most αSyn positive cases initial brain lesions appear in the lower brainstem areas (caudo-rostral), while others exhibit an early involvement of the amygdala (amygdala-predominant).^12–14^

Amygdala-predominant αSyn pathology is often associated with co-morbid AD neuropathologic changes (ADNC),^7,13–15^ which includes amyloid β (Aβ) plaques, neuritic plaques, and neurofibrillary tangles (NFT) consisting of abnormal phosphorylated-tau protein (pTau). Within the spectrum of AD, various subtypes have been identified based on the severity of NFT pathology including limbic-predominant, hippocampal sparing, and typical AD.^7^ Among these, the limbic-predominant subtype is characterized by more severe limbic system pathology compared to that in the neocortex^16^ and has been recently identified to have the highest occurrence of amygdala-predominant αSyn pathology.^3^

Hippocampal degeneration is one of the hallmark of AD, affecting about 80% of AD patients.^17^ The limbic-predominant subtype of AD was shown to be characterized by the most severe volume loss of that region.^18^ Both the APOE ε4 variant^19^ and pTau pathology^20^ have been linked with increased hippocampal atrophy and recent studies also highlight the significant impact of TDP-43.^21,22^ Furthermore, cell loss in the CA1 region has been linked to arteriolosclerosis,^23^ cerebral amyloid angiopathy (CAA),^24,25^ epilepsy,^26^ as well as metabolic imbalances such as ischemia and hypoglycemia.^27^ This demonstrates that hippocampal degeneration is a multifaceted phenomenon, potentially influenced by different sets of factors in different individuals.

In patients with LBD, αSyn pathology primarily affects the CA2 region.^28^ Despite the severity of αSyn pathology in CA1 correlating with memory impairment in LBD,^29^ studies investigating the CA1 and subiculum regions have found no clear effects of αSyn pathology on degeneration of these regions.^30,31^ It is important to note that patients with αSyn aggregates often have an abundance of other misfolded proteins.^32^ Additionally, αSyn pathology has been observed to interact with pTau,^33,34^ Aβ,^35,36^ and TDP-43^37–39^ pathology, which adds complexity to understanding its impact on neuronal loss in the hippocampus.

In this study, we investigated whether αSyn pathology, similarly to other AD-related pathologies, influences the degeneration of the CA1 region of the hippocampus. We categorized αSyn pathology into amygdala-predominant and caudo-rostral variants, hypothesizing that the impact on hippocampal degeneration might vary based on the origin of the protein accumulation. We then evaluated the differences in neuropathological profiles of AD patients with these two patterns of αSyn pathology distribution. Finally, we used structural equation modeling to explore how pTau, TDP-43, and αSyn interact within the limbic system, and which of these pathologies is directly related to neuron loss in the CA1 region.

## Materials and methods

### Human samples

We examined 291 autopsy brains obtained from university or municipal hospitals in Leuven (Belgium, Research Ethics Committee (EC) UZ/KU Leuven identifiers: S52791, S55312, S59292, S64363), Bonn, Offenbach am Main, and Ulm (Germany, EC UZ/KU Leuven identifier: S59295, Ulm identifier: 58/08) and GE Healthcare (ClinicalTrials.gov identifiers NCT01165554 & NCT02090855). All of the experiments have been performed after ethical clearance by UZ/KU-Leuven ethical committee (S65147). The brain samples were collected in accordance with local legislation and the federal laws governing the use of human tissue for research in these three countries. Clinical files have been analyzed for the presence of parkinsonism symptoms, diagnosis of epilepsy and dementia status in our cohort. The degree of dementia at the time of death was determined retrospectively using an estimate of the Clinical Dementia Rating (CDR) global score^40^ and/or Mini Mental Score Examination (MMSE).^41^ The MMSE has been converted to a CDR score using published cut-off values.^42^ A description of the neuropathological and clinical characteristics of the study group can be found in Supplementary Table 1.

Exclusion criteria listed in Supplementary Table 2 were rigorously applied to ensure the homogeneity of the study population. While cases with a clinical diagnosis of seizure disorder were not excluded, this information was utilized as a confounding factor in our primary analysis. In cases featuring TDP-43 proteinopathies, only those with clear features of either Amyotrophic Lateral Sclerosis or FTLD-TDP subtypes (diagnosed according to the criteria outlined by)^43^ were excluded, owing to the common co-occurrence of pTDP-43 pathology in Alzheimer’s disease.^22,44^

### Immunohistochemistry

We used formalin-fixed tissue samples from the left brain hemisphere (for *n* = 235) and right (for *n* = 44), including the anterior medial temporal lobe (MTL), posterior MTL with hippocampus, middle frontal gyrus, occipital cortex, midbrain, pons, and medulla. The tissue was embedded in paraffin and sectioned into 5μm thick sections using a microtome (Thermo Fisher Scientific).

Immunohistochemistry was employed to stain these sections. Detailed information about the antibodies used and the regions stained can be found in Supplementary Table 3. A robotic autostainer (Leica Microsystems) was used to perform tissue deparaffinization. Antigen retrieval was carried out in a PT Link Module (Dako) using EnVision Flex Target Retrieval Solution Low pH (a citrate-based buffered solution with a pH of 6.1, Dako). An additional step involving a 5-minute incubation with formic acid was included for αSyn staining. To block endogenous peroxidase activity, EnVision FLEX Peroxidase-Blocking Reagent was applied to the tissue for 5 minutes. Tissue sections were incubated overnight in a humid chamber with a reagent containing the primary antibody. The next day, anti-mouse Horseradish peroxidase (HRP)-linked secondary antibodies were used for staining against pTau and αSyn, while the anti-rabbit ABC-HRP Kit (VECTASTAIN) was employed for pTDP-43 staining to enhance the signal. Finally, 3,3’-diaminobenzidine solution (Liquid DAB + Substrate Chromogen System, Dako) was used to visualize the binding between the primary antibody and secondary antibody. All slides were counterstained with hematoxylin using an autostainer, and mounting was performed with an automated cover-slipper (Leica Microsystems). A ZEISS Axio Imager 2 Microscope equipped with Camera Axiocam 506 and DM2000 LED Leica microscope equipped with a Leica DFC7000 T digital camera ware used for the examination and digital photography of the tissue.

### Pathology assessment and quantification

Pathological assessment was conducted for all cases following established protocols. This included evaluating Braak stages of neurofibrillary tangles progression,^45^ phases of Aβ deposition in MTL,^46^ CERAD measurements of neuritic plaques.^47^ The Braak NFT stages, Aβ MTL phases, and CERAD measurements were translated into A, B, and C-scores, respectively.^48^ An "ABC score" reflecting the severity of ADNC was calculated according to the National Institute on the Alzheimer’s Association guidelines.^48^

Cases that exhibited αSyn immunoreactivity in the medulla or amygdala underwent further screening for αSyn pathology in the pons, midbrain, posterior hippocampus, posterior temporal cortex, and frontal cortex. The Braak LBD Stage, a widely recognized staging system for Parkinson’s disease αSyn pathology,^9^ was calculated. For the evaluation of αSyn lesion severity, we applied well-established semi-quantitative criteria as previously outlined^9^ and the third report from the Dementia with Lewy Bodies (DLB) Consortium^11^ for each of these regions, except for the locus coeruleus in the pons as the tissue was not always available. Our assessment covered all types of αSyn lesions, including Lewy neurites, Lewy bodies, and glial inclusions within gray matter. The representative levels for each severity category in each brain region are depicted in Supplementary Fig. 1. We excluded 7 cases with missing data on αSyn severity in more than two regions leaving a sample of *n* = 105 αSyn positive cases, and imputed values from 14 cases with a single missing data point (2.2% of all values). We calculated the global burden score for αSyn by summing the severity scores across the six brain regions. Additionally, we computed the ratio of αSyn pathology severity scores as follow: (*amygdala + posterior temporal lobe*) */* (*medulla + midbrain*). We used this ratio to stratify patients into two αSyn pathology spreading patterns, considering cases with a ratio higher than one as ’amygdala-predominant’ and the rest as ’caudo-rostral.’

Manual quantification of neurons in the CA1 region was performed on hematoxylin-stained slides of the posterior hippocampus. Three non-overlapping photographs were taken, each containing only the CA1 region, with a magnification of x200. Neurons were identified based on their morphology using QuPath^49^ software, and neuronal density was calculated per mm^2^. Selection criteria for neurons included the presence of a spherical nucleus and a visible, often pyramid-shaped, cytoplasm.

We defined a group of symptomatic AD cases as having a clinical diagnosis of dementia with moderate or severe ADNC neuropathological changes (“ABC” score 2 or 3). For this group, we additionally assessed the severity of pTau pathology in CA1 by taking three photographs of this region from AT8 stained slides and quantifying the proportion of pTau-positive neurons to the total number of neurons. Additionally, we screened for the presence of pTDP-43 lesions in the amygdala, posterior hippocampus, and frontal cortex, and for cases with all three regions available, we classified the LATE-NC stages using the updated guidelines.^50^ We also investigated the positivity of pTDP-43 lesions in the dentate gyrus of the posterior hippocampus. Finally, we assessed the type (type 1 with capillary involvement vs. non-capillary type 2) and severity of CAA^51,52^.

### Data analysis

Data analysis was conducted using R Studio software (version 2023.06.0 with R-4.3.1). An a priori power analysis was conducted using G*Power version 3.1.9.7^53^ For multiple analyses, we calculated the a priori sample size for our largest model (8 predictors) to achieve 95% power with α = 0.05, detecting a medium effect for R2 deviating from zero. The analysis indicated a minimum sample size of 160, which is less than the 262 cases we utilized. To determine the minimum sample size required to test the study hypothesis of differences between three groups of AD patients, results indicated that the required sample size to achieve 95% power for detecting a large effect (f = 0.4), at a significance criterion of α = 0.05, was *n* = 100 for ANOVA very close to sample size used by our study (*n* = 99).

To determine the significant relationships between two categorical variables, we employed a two-sided Fisher’s exact test. For variables with more than two levels that reach the statistical level of 0.05, the two-sided pairwise Fisher’s exact tests were then conducted. For examining differences in continuous variables between more than two groups, we assessed the normality assumption in each group using a Shapiro-Wilk test. In cases where this test was not satisfied, as well as for ordinal variables, we applied a Kruskal–Wallis test with a Dunn-Bonferroni post-hoc test. For normally distributed variables, we used an ANOVA with the Dunn-Bonferroni post-hoc test. For comparing not-normally distributed values between two groups we used two-sided Wilcoxon Rank Sum and Signed Rank Tests.

Multiple linear regression was employed to examine the relationship between several independent variables and neuronal density in CA1 region. To validate our results, we tested whether our models meet the necessary assumptions. We visually inspected scatter plots to ascertain whether continuous and ordinal independent variables exhibited a linear relationship with the dependent one. The normality of residuals was examined using the Shapiro-Wilk test, and dependency of residuals on the values of the independent variables was assessed with the Breusch-Pagan test for heteroscedasticity. Additionally, we calculated the Variance Inflation Factor for each predictor to ensure the absence of multicollinearity.

For the correlation analysis between αSyn severity in various brain regions and neuronal density, we employed the semi-partial Spearman correlation function while controlling for age at the time of death using the ppcor library.^54^ In cases with a single missing value, we assumed, based on the assumption of spreading, that the αSyn severity for each patient forms a sequence. As such, missing values for the midbrain, amygdala, posterior hippocampus, and temporal cortex were imputed using the na.approx function from the zoo library.^55^ To mitigate the false discovery rate, P-values obtained from correlation analysis, Wilcoxon tests, and all post-hoc tests were adjusted using the Benjamini–Hochberg method.

Finally, using only data from AD patients, we modeled the association between observable neuropathological parameters through path analysis within the structural equation modeling (SEM) framework (Supplementary Fig. 3, lavaan package version 0.6.16).^56^ SEM analysis facilitates the examination and quantification of both direct and indirect relationships among numerous variables,^57^ a method previously applied successfully in the study of multifactorial diseases such as diabetes.^58^ We posited that the amygdala-predominant αSyn pathology exert a downstream effect on other variables, thereby treating it as the sole exogenous variable. The endogenous variables in our model included the percentage of neurons affected by pTau in the CA1 region, LATE-NC stages, and neuronal density in the CA1 area. The model’s parameters were estimated using the Maximum Likelihood method, employing the standard NLMINB optimization technique. To verify the model’s accuracy, we evaluated its fit using both relative indices (e.g., Comparative Fit Index [CFI], Tucker-Lewis Index [TLI]) and absolute indices (e.g., Standardized Root Mean Square Residual [SRMR], Root Mean Square Error of Approximation [RMSEA]).

### Data availability

Summary statistics of basic demographic, clinical, and neuropathological parameters for each group used in this study are available in the Supplementary Table 1. Due to legislation and privacy protection any medical reports and files of the cases included in this study cannot be made available. Pseudonymized data will be provided by the corresponding author upon reasonable request.

## Results

### α-synuclein pathology is associated with neuronal loss in the hippocampal CA1 region

Using a cohort of 291 both demented and non-demented individuals with varying degrees of neuropathological changes we investigated the relationship between the presence of αSyn pathology and neuronal density in the CA1 subfield of the hippocampus. To account for the expected effect of Aβ and Tau pathology on neuronal loss, we stratified cases based on the overall severity of ADNC. We operationalized αSyn-positive cases as exhibiting any degree of pathology in investigated regions.

We observed that neuronal densities in individuals with αSyn positivity were lower (*mean*(*SD*) = 138.4(58.4)) compared to those without αSyn pathology (*mean*(*SD*) = 166.1(59.6), Fig. 1A). A Wilcoxon test confirmed significant differences between these two groups (*Z* = 24.78, *P* < 0.001) indicating higher CA1 neuronal loss in individuals with αSyn burden. Next, we investigated whether ADNC severity contributed to the observed effects of αSyn pathology. Fig. 1B presents a violin plot illustrating CA1 neuronal density for αSyn-positive and αSyn-negative groups, categorized by the severity of ADNC scores. Interestingly, the lowest neuronal density was evident in cases exhibiting both αSyn pathology and severe ADNC (*mean*(*SD*) = 104.5(53.64)).

**Figure 1.**
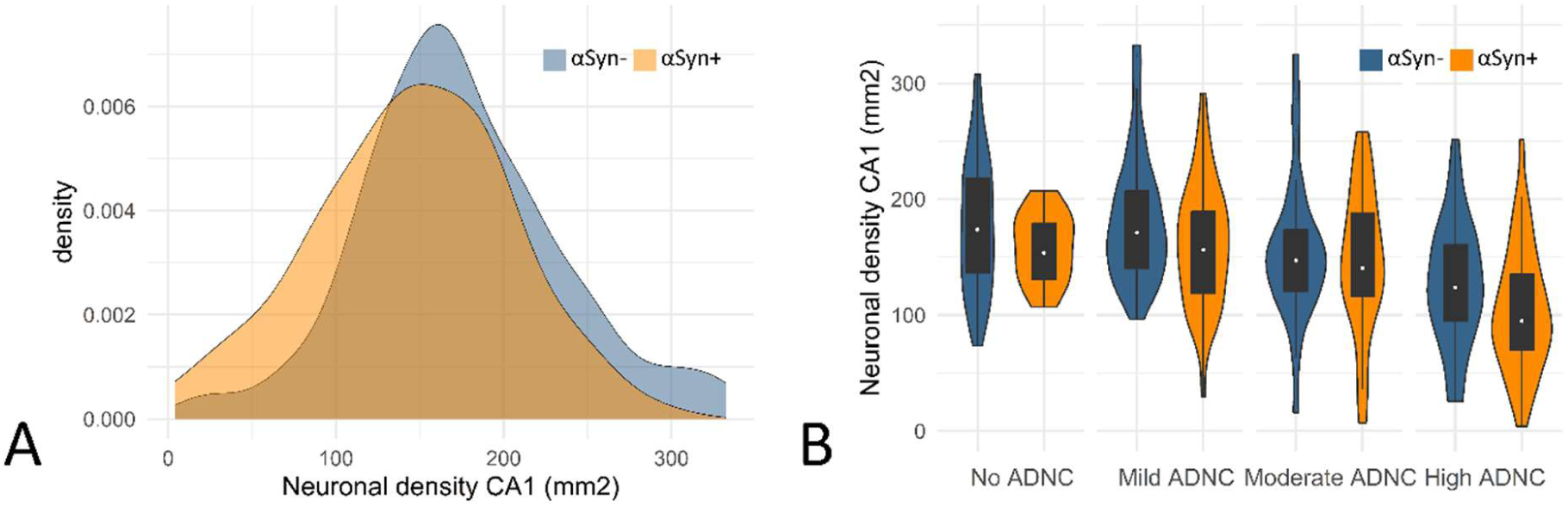
Relationship between hippocampal neuronal density, the presence of αSyn pathology and ADNC severity. **(A)** The distribution of neuronal density per 1 mm² in the CA1 region of the posterior hippocampus is depicted for αSyn-negative cases (αSyn-, *n* = 173) and αSyn-positive cases (αSyn+, *n* = 105). **(B)** A violin plot illustrates the differences in CA1 neuronal density within the same population, stratified by the presence of αSyn, and categorized by the severity of ADNC on the x-axis. Each boxplot within the violin plot represents the 1st to 3rd interquartile range, with the white point indicating the median.

To examine the degree of how much each pathology contributed to hippocampal degeneration we developed a multiple regression model (*n* = 290, Supplementary Table 4-5 [Model 1], Supplementary Fig. 2) with presence of αSyn pathology, Braak NFT stages, Aβ MTL phases, sex, and age at death as predictors for CA1 neuronal density. We showed that the presence of αSyn pathology was a significant independent predictor of CA1 neuronal density (*β* = -18, *P* = 0.01), even after accounting for the effects of Braak NFT stages (*β* = -7.1, *P* = 0.005) and Aβ (*β* = -6.9, *P* = 0.022). Age at death (*β* = 0.1, *P* = 0.749) and sex (*β* = 10.2, *P* = 0.127) did not demonstrate a significant impact on CA1 neuronal density in this model.

### Amygdala-predominant α-synuclein pathology pattern is common in brains with Alzheimer’s disease

Next, we decided to investigate the hypothesis that the amygdala-predominant variant of αSyn pathology contributes to limbic degeneration. To address this, we assessed the αSyn pathology severity in six brain regions from 112 cases exhibiting varying levels of αSyn (Braak PD stage 1-6). The distribution of obtained scores for each region is shown in Supplementary Table 6. For these αSyn positive cases, we calculated the ratio between αSyn pathology severity in the MTL (amygdala+temporal cortex) and brainstem (medulla+midbrain) as well as the αSyn pathology global burden score.

In our sample, the overall severity of αSyn pathology, reflected by the global burden score, was comparable between groups (Fig. 2A). Although it was the lowest in cases without any ADNC pathology (*mean*(*SD*) = 6.2(4.5)), Kruskal-Wallis test showed no group effect (*H*(3) = 2.6, *P* = 0.46, Supplementary Table 6). On the other hand, we observed a trend towards increased MTL to brainstem αSyn pathology severity ratio with increasing ADNC (Fig. 2B). The group exhibiting the most severe ADNC displayed the highest observed αSyn pathology ratio (*median* = 2), while the group without ADNC (*median* = 0.33) had no case surpassing a value of 1, suggesting that all these cases exhibited the caudo-rostral variant. Kruskal-Wallis test confirmed the differences between groups (*H*(3) = 28.95, *P* = <0.001). Dunn’s multiple comparisons test showed significant differences between cases with the highest ADNC severity and all other groups (Supplementary Table 7). Cases with a ratio higher than one were classified as amygdala-predominant’ (*n* = 34), and the rest were classified as ’caudo-rostral’ (*n* = 71).

**Figure 2.**
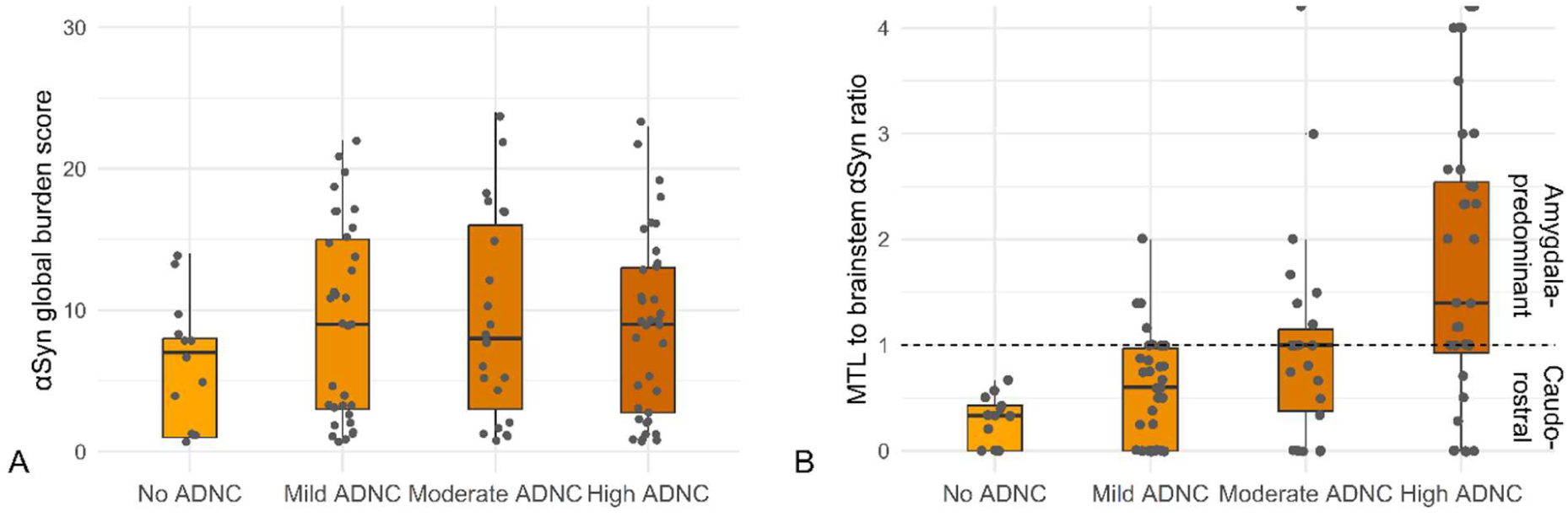
Changes in overall αSyn pathology burden and prevalence of amygdala-predominant variant with increasing ADNC levels. Boxplots with scattered points depicting the mean αSyn pathology global burden score **(A)** and the ratio of αSyn pathology severity in MTL to brainstem **(B)** for different levels of ADNC. Cases with higher pathology severity in MTL than in the brainstem (ratio > 1) have been labeled as having an ‘amygdala-predominant’ pattern, and other cases as ‘caudo-rostral’. Ratio values that reach infinity due to no pathology in the brainstem are indicated as half-dots. The horizontal bold lines represent median values, the box margins illustrate the 25% and 75% quartiles, while the whiskers display the lower and upper range values.

We then tested whether an amygdala-predominant pattern was driving the association with CA1 neuronal density by adding two spreading patterns as predictors to the regression model (*n* = 283, Supplementary Table 4-5 [Model 2]). We demonstrated that the amygdala-predominant αSyn pathology pattern was the primary predictor of neuronal density in the CA1 region in our sample (*β* = -39.6, *P* < 0.001). Caudo-rostral αSyn pathology was not significantly negatively associated with CA1 degeneration (*β* = -5.41, *P* = 0.48). Finally, to exclude other potential factors impacting neuronal density in our sample, we extended Model 2 with additional predictors including postmortem interval (PMI), clinical diagnosis of epilepsy, and the hemisphere from where the tissue had been taken (Supplementary Table 4-5 [Model 3]). No impact of PMI (*n* = 262, *β* = 0.2, *P* = 0.134) and hemisphere (*β* = -1.9, *P* = 0.847) was observed. As expected, epilepsy was negatively associated with CA1 neuronal density (*β* = -34, *P* = 0.023). The association between the amygdala-predominant αSyn pathology pattern and CA1 degeneration persisted (*β* = -42.8, *P* < 0.001).

### The severity of α-synuclein pathology in amygdala, but not CA1, correlates with CA1 neuronal density

To investigate whether the effect of αSyn pathology on CA1 degeneration resulted from its general presence in the brain or its intensity in particular areas, we conducted a semi-partial Spearman correlation analysis. αSyn pathology severity scores in various brain regions were considered the predictor variables, neuronal density in the CA1 region the outcome variable, with age at death as covariate. Only αSyn-positive cases with complete information about severity across brain regions were included in this analysis (*n* = 105). The results of these correlations are presented in Fig. 3, and the associated corrected P-values are detailed in Supplementary Table 8.

**Figure 3.**
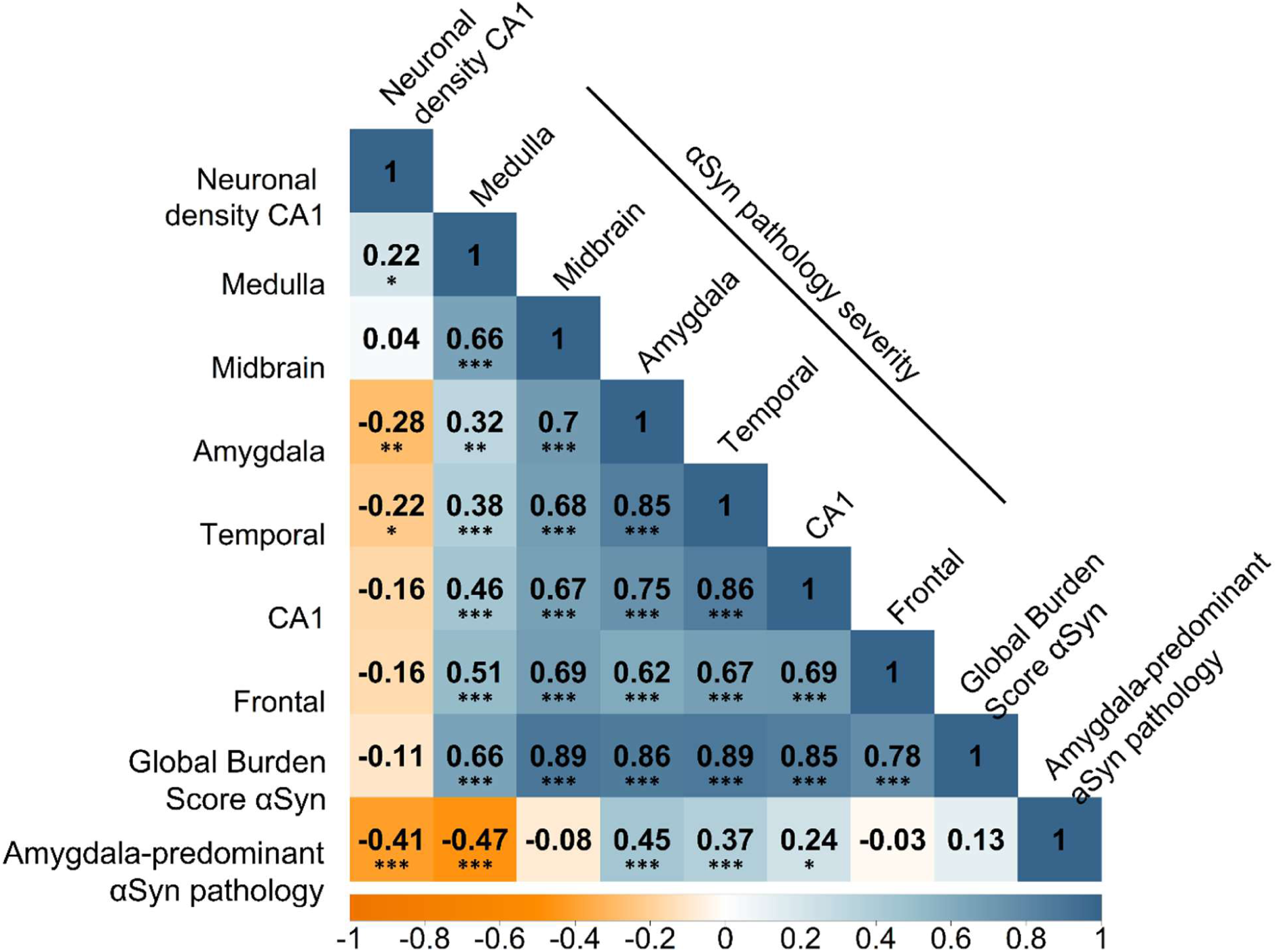
Association between αSyn pathology severity if different brain regions and hippocampal damage. This figure illustrates the results of correlation analyses assessing the relationship between neuronal density in the hippocampal CA1 region and various measures of αSyn pathology. The investigated parameters include αSyn pathology severity in different brain regions, global burden score of αSyn pathology, and the presence of an amygdala-predominant αSyn pathology spreading pattern. The pattern of spreading originating in the amygdala, along with the severity of αSyn pathology in the amygdala, demonstrated the most robust association with decreased neuronal density in the CA1 region. Each cell in the matrix displays Spearman semi-partial correlation coefficients (rho) death representing the strength and direction of the correlation between the corresponding pairs of values while controlling for age at death. Medulla – medulla oblongata, temporal – temporal cortex, frontal – frontal cortex; * <0.05, ** <0.01, ***<0.001

This analysis confirmed that neuronal density of the hippocampal CA1 region is negatively associated to severity of αSyn pathology in amygdala (*ρ*=-0.29, *P* = 0.005), posterior temporal cortex (*ρ*=-0.22, *P* = 0.03), and presence of amygdala-predominant αSyn pathology spreading variant (*ρ*=-0.42, *P* = <0.001). Interestingly, no significant relation between CA1 neuronal density and severity of αSyn pathology in CA1 was observed (*ρ*=-0.17, *P* = 0.121) nor in the frontal cortex (*ρ*=-0.16, *P* = 0.140). As suspected, no correlation was found between αSyn pathology in the substantia nigra of the midbrain and neuronal density in the CA1 region (*ρ*=0.03, *P* = 0.888). Lastly, a small positive effect was observed for the medulla (*ρ*=0.22, *P* = 0.039) likely due to its high inverse correlation with the amygdala-predominant αSyn pathology variant (*ρ*=-0.47, *P* = <0.001).

Furthermore, all severity scores for αSyn pathology exhibited positive intercorrelations. The strongest relation was observed between the amygdala and temporal cortex (*ρ*=0.84, *P* = <0.001) and between CA1 and temporal cortex (*ρ*=0.87, *P* = <0.001), whereas the weakest correlation was between amygdala and medulla αSyn pathology severity (*ρ*=0.32, *P* = 0.002). Despite these strong associations between αSyn pathology severities for different regions, the global burden score did not correlate with decreased CA1 neuronal density (*ρ*=-0.08, *P* = 0.508).

### Diverse neuropathological profiles in Alzheimer’s patients with two α-Synuclein patterns

Next, we sought to investigate whether AD cases with amygdala-predominant and caudo-rostral αSyn pathology patterns differ in their spectrum of co-pathologies. We hypothesized that the increased neuronal degeneration in the amygdala-predominant pattern may be associated with an accumulation of other misfolded proteins. To explore this, we identified in our cohort 99 cases with clinically confirmed dementia exhibiting moderate to severe ADNC. For this analysis, we used additional information on the percentage of neurons affected by pTau in CA1, the LATE-NC stages, the presence of pTDP-43 in the dentate gyrus, as well as the severity and type of CAA. Descriptive statistics and the number of cases for each parameter are available in Supplementary Table 9.

Within our cohort of AD patients, 48 had no αSyn pathology, 29 exhibited the amygdala-predominant variant, and 22 showed the caudo-rostral pattern. We examined differences in neuropathological and demographic characteristics between these groups (Fig. 3) using ANOVA or Kruskal–Wallis tests (Supplementary Table 10) followed by Dunn-Bonferroni post-hoc analysis (Supplementary Table 11), and pairwise Fisher’s test (Supplementary Table 12).

We observed that the selected groups did not differ in their age at death (Fig. 4A, *F*(2,96) = 0.66, *P* = 0.518). Regarding sex proportions, we observed a small but not significant trend between groups, with the AD αSyn negative group having the highest proportion of women and AD with the caudo-rostral αSyn pathology pattern having the highest proportion of men (Fig. 4B, *P* = 0.230). Overall, groups did not differ significantly in their basic demographic characteristics, justifying the comparison of their other neuropathological parameters. Additionally, both groups of AD patients with αSyn pathology had the same levels of global burden score reflecting the overall severity of the pathology (Fig. 4C, *Z* = 0, *P* = 1). As predicted, AD with amygdala-predominant αSyn pathology had the lower CA1 neuronal density than AD and AD with caudo-rostral pattern (Fig. 4D, *mean*(*SD*) = 99.47(52.04)), but the difference was not statistically significant after adjustment for multiple comparisons (*Z* = -2.2, *P* = 0.084, *Z* = -1.87, *P* = 0.092).

**Figure 4.**
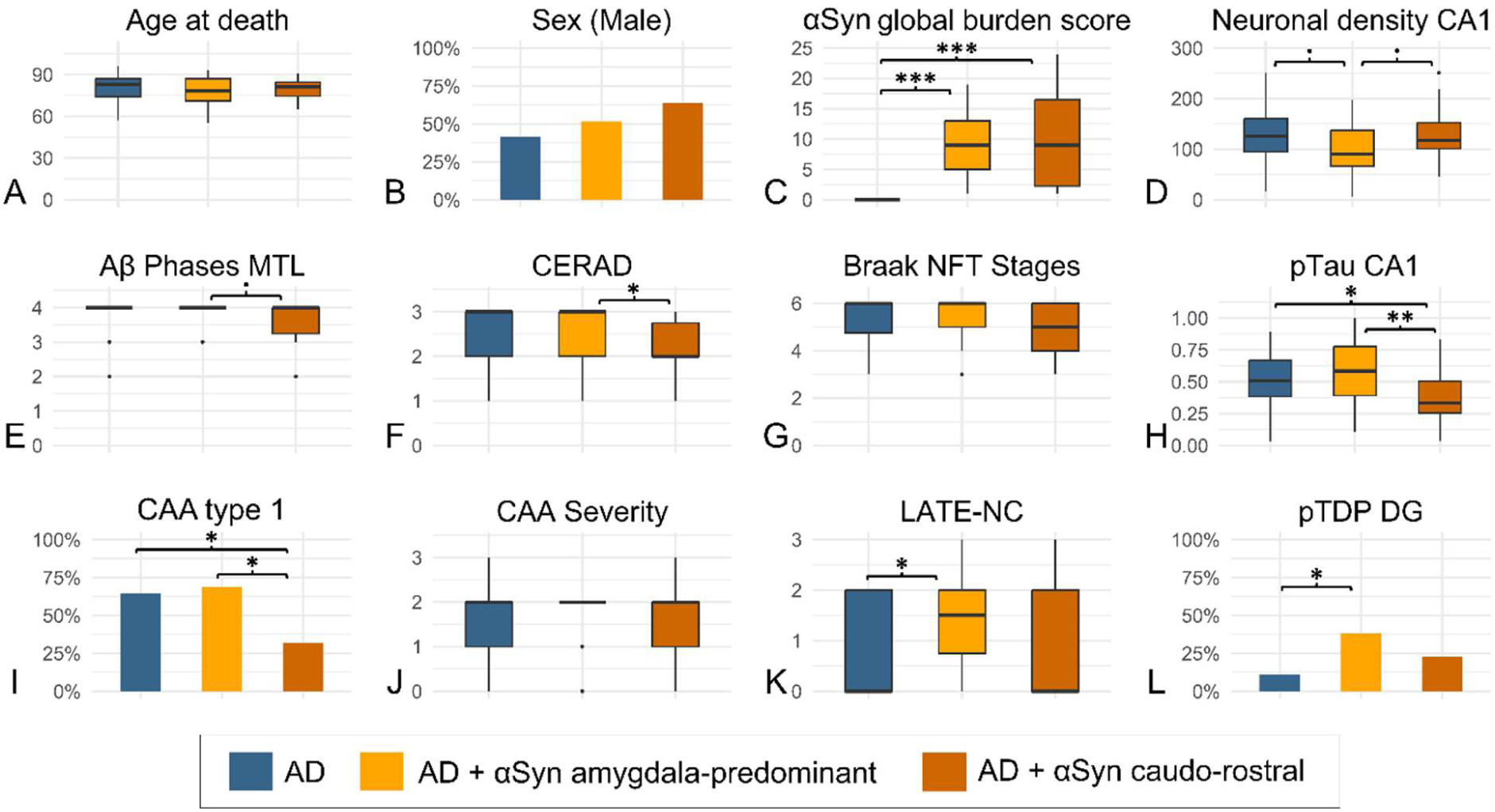
Characteristics of AD patients with and without αSyn pathology. AD patients (*n* = 99) have been stratified based on the presence and distribution of αSyn pathology into an αSyn-negative group (AD) and two αSyn-positive groups with amygdala-predominant or caudo-rostral variants. **(A-J)** Boxplots with horizontal bold lines indicate median values, the box margins represent the 25% and 75% quartiles, while the whiskers show the lower and upper range values. **(K-P)** Barplots for binary variables show the percentage of cases positive for the clinical or neuropathological feature; .<0.1, *<0.05, **<0.01, ***<0.001

While patients with the amygdala-predominant αSyn pathology pattern displayed comparable levels of ADNC to αSyn-negative individuals, those with the caudo-rostral variant generally exhibited a lower degree of ADNC severity (Fig. 4E-H). Statistical analysis revealed that patients with the caudo-rostral αSyn pathology pattern patients significantly differed from the αSyn-negative group in the severity of neuritic plaques, reflected by the CERAD score (*Z* = 2.66, *P* = 0.023) and that they had fewer pTau-positive neurons in CA1 than the other groups (*Z* = 2.24, *P* = 0.038; *Z* = 3.16, *P* = 0.005). Interestingly, patients with caudo-rostral αSyn pathology had the lowest rate of developing type 1 CAA (Fig. 4I, *P* = 0.0285), although there was no difference in CAA severity among groups (Fig. 4J, *H*(2) = 2.1, *P* = 0.351). On the other hand, AD with the amygdala-predominant variant had the same prevalence of CAA type 1 as αSyn-negative patients (*P* = 0.81). However, AD patients with amygdala-predominant αSyn pathology had significantly higher LATE-NC stages (Fig. 4K, *Z* = 2.84, *P* = 0.013) and showed more prevalent pTDP-43 inclusions in dentate gyrus (Fig. 4L, *P* = 0.02).

### TDP-43 and Tau mediate amygdala the impact of α-Synuclein on hippocampal degeneration

To fully understand the interactions between AD-related neuropathologies in the MTL and CA1 degeneration, we constructed a path analysis model (Supplementary Fig. 3). The model estimated 8 parameters using data from 90 out of the total 99 observations from AD. The remaining 9 cases had missing values for at least one of the neuropathological parameters used. The model demonstrated a good fit based on both relative and absolute fit measures, adhering to standard guidelines (Supplementary Table 13)^59^. The structure of this model and the standardized estimates are depicted in Figure 5A and additional model details can be found in Supplementary Table 13.

**Figure 5.**
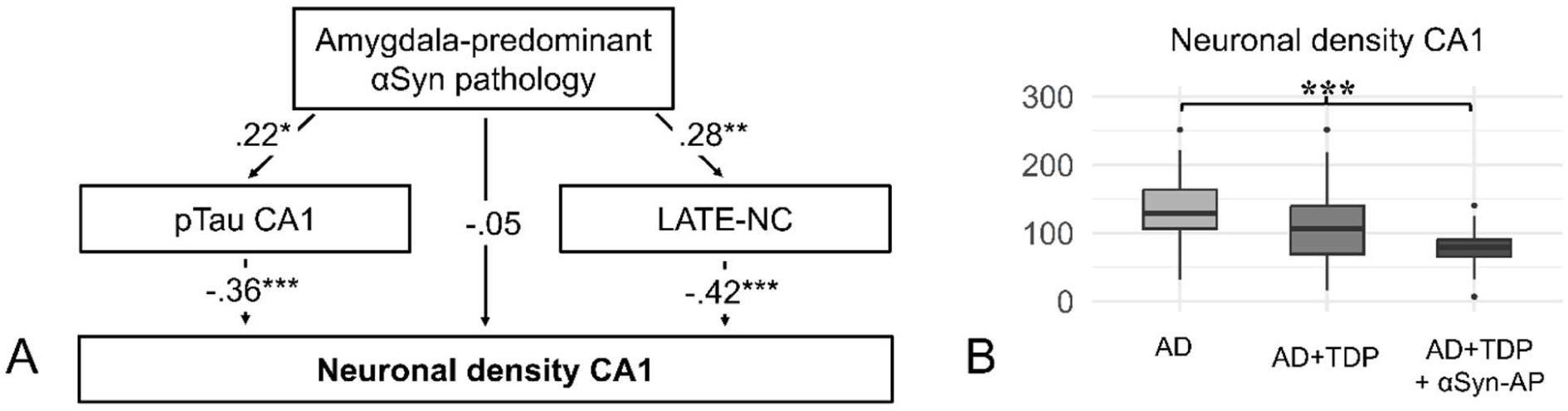
The interactions between AD-related neuropathological changes and decreased CA1 neuronal density. **(A)** The graphical representation of outcomes from a Structural Equation Model designed to test whether amygdala-predominant αSyn pathology has direct association with neuronal density in CA1 or whether it is mediated by other pathologies in AD patients (*n* = 90). The observed variables (neuropathological parameters) are represented by square boxes and include the presence of amygdala-predominant αSyn pathology, the percentage of neurons affected by pTau in CA1, stages of LATE-NC and neuronal density in CA1 region. Arrows represent regression paths connecting the variables, with the direction of the arrow indicating the flow from the independent to the dependent variable. These arrows are annotated with standardized regression coefficients (estimates). Symbols next to the coefficients indicate the levels of statistical significance represented by P-values. **(B)** Boxplot showing differences between AD patients without hippocampal TDP-43 pathology (AD, *n* = 59), AD with hippocampal TDP-43 (AD+TDP, *n* = 25), and AD patients with both hippocampal TDP-43 and amygdala-predominant variant of αSyn pathology (AD+TDP+αSyn-AP, *n* = 12). Horizontal bold lines indicate median values, the box margins represent the 25% and 75% quartiles, while the whiskers show the lower and upper range values. * <0.05, ** <0.01, *** <0.001

Based on the model results we observed that the presence of the amygdala-predominant αSyn pathology pattern was related to both an increase in pTau pathology density in CA1 (*estimate*[*SE*] = 0.11 [0.05], *P* = 0.032) and LATE-NC (*estimate[SE]* = 0.62 [0.28], *P* = 0.006). We also observed the significant effect of pTau (*estimate[SE]* = -81.14 [19.9], *P* < 0.001) and LATE-NC (*estimate[SE]* = -21.261 [4.47], *P* < 0.001) on CA1 neuronal density. However, we could not see any direct effect of amygdala-predominant αSyn pathology (*estimate[SE]* = -5.32 [10.32], *P* = 0.515).

To further illustrate the significant indirect impact of αSyn pathology on CA1 degeneration, we classified AD patients into three group: those without hippocampal pTDP-43 (*n* = 60), those with hippocampal pTDP-43 but without amygdala-predominant αSyn pathology (*n* = 23), and those with both hippocampal pTDP-43 and amygdala-predominant αSyn pathologies (*n* = 13). Three cases could not be classified because of missing values. Our analysis revealed that AD patients with all three pathologies exhibited the lowest neuronal density (*mean* (*SD*) = 75.3 (36.2), Fig. 5B), followed by the group with only TDP (*mean* (*SD*) = 108 (60.1)), and the highest in the group without these two pathologies (*mean* (*SD*) = 133 (47.9)). A Kruskal-Wallis test with post-hoc comparisons revealed). Our data followed a trend, with each additional co-pathology leading to higher CA1 neuronal loss. Nevertheless, differences between AD patients with and without pTDP-43 (Z = 2.06, P = 0.06), as well as those with pTDP-43 alone and those with the triple pathology (Z = 1.81, P = 0.07), were not statistically significant. However, we observed a significant difference between AD patients with all three pathologies compared to the group without (Z = 3.7, P < 0.001).

## Discussion

In this study, we analyzed a cohort with varying degrees of ADNC and αSyn pathology, uncovering a relationship between αSyn pathology and neuronal loss in the hippocampal CA1 region. Further analysis indicated that this correlation is more pronounced in cases with an amygdala-predominant variant, and that the severity of αSyn pathology in the amygdala, but not CA1, more strongly correlates with neurodegeneration. This finding implies that the primary determinant of cell loss in CA1 may not be determined by an increase in αSyn pathological burden in this region. Consequently, we explored alternative lesions that might mediate this relationship, uncovering distinct neuropathological profiles between AD cases with caudo-rostral and amygdala-predominant αSyn pathology variants. Employing a structural equation model, we demonstrated that in AD patients the influence of αSyn pathology on CA1 neuronal loss is mediated by the severity of pTau and TDP-43 pathology in the limbic system, both directly associated with CA1 degeneration.

A key discovery of our study is the significant role of the amygdala-predominant αSyn pathology in contributing to limbic system neurodegeneration. Previous studies have not found a clear association between amygdala-predominant αSyn pathology and cognitive impairment,^7,60^ potentially due to the subtyping method, which categorized cases as amygdala-predominant only if lesions were limited to that region and therefore mild. To address this limitation, we employed a ratio of αSyn pathology severity in the limbic system to the brainstem. Our analysis revealed that AD patients with amygdala-predominant αSyn and TDP-43 pathology exhibit the lowest neuronal density in CA1. This finding aligns with studies demonstrating that AD patients with TDP-43 and αSyn co-pathology experience the most severe cognitive decline.^61^ Furthermore, our results support a recent study that identified a high prevalence of amygdala-predominant αSyn and LATE-NC in the limbic-predominant AD subtype,^3^ which suggests that such patients may experience less widespread but more severe damage originating in the limbic system.

The amygdala is recognized as a critical region for the accumulation of neurodegenerative pathologies, including pTau, TDP-43, and αSyn.^62^ It is tempting to speculate that close proximity of pathological aggregates may facilitate interactions among them, potentially amplifying pathology, which can then spread to adjacent brain regions (Fig. 6B). Previous studies have demonstrated that αSyn aggregates induce the fibrilization of Tau^34^ and its aggregation into NFT-like structures.^63^ TDP-43 pathology, known to exacerbate neuronal loss and contribute to hippocampal sclerosis in AD^22,44^ and LBD,^64^ has also been shown to interact with αSyn pathology.^37–39^ In vitro studies showed the formation of hybrid fibrils between these proteins,^65–67^ amplifying their neurotoxic effects. Moreover, neuronal degeneration has been associated with the interaction between Aβ, pTau, and αSyn with cellular prion protein in cell cultures.^68^ Finally, there is a possibility that the presence of αSyn fibrils in CA1 pathology can contribute to neuronal loss, but our analysis was not sensitive enough to detect it. In LBD, αSyn accumulation typically leads to the loss of dopaminergic neurons in the brainstem, with various proposed mechanisms for its neurotoxic effects, including disruption of autophagy.^69,70^ Thus, the neurodegenerative impact of αSyn might results, at least partially, from its ability to enhance Tau and TDP-43 pathology, form more toxic fibrils with TDP-43, or directly disrupt cellular processes.

**Figure 6.**
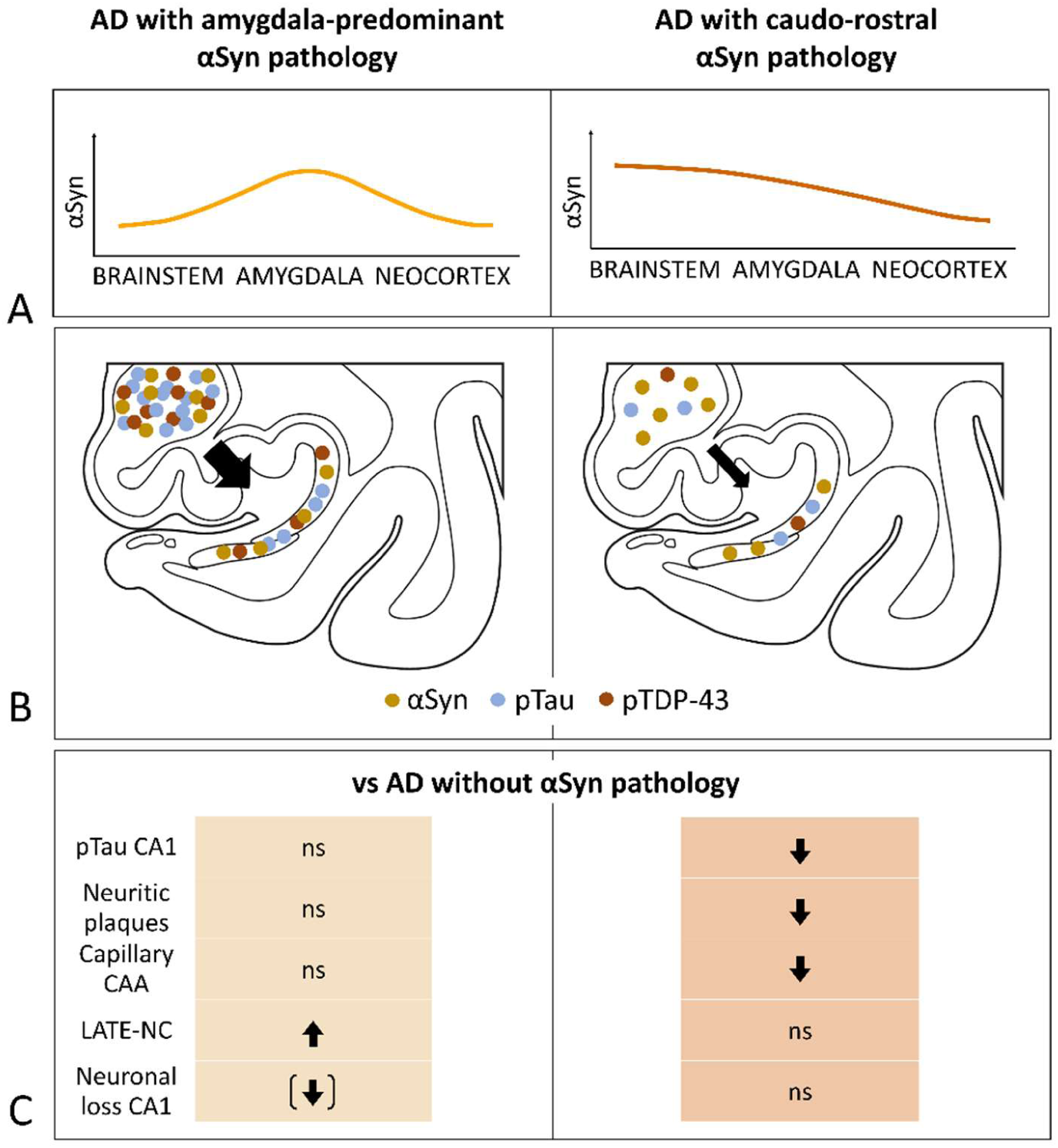
A graphical abstract illustrating the distinctions between AD with caudo-rostral and amygdala-predominant αSyn pathology. **(A)** Depiction of the simplified distribution of αSyn pathology severity for two spreading patterns. **(B)** Proposed mechanisms underlying increased hippocampal neuronal loss in AD with amygdala-predominant αSyn pathology, where the amygdala acts as a hub for interaction and heightened aggregation of αSyn, pTDP-43, and pTau, resulting in elevated triple pathology in the hippocampus. The interaction is less pronounced in the caudo-rostral variant, likely due to a shorter incubation time for all misfolded proteins. **(C)** Summary panel presenting the study’s findings comparing AD with two αSyn pathology spreading patterns to AD without αSyn pathology. An upward arrow indicates an increase in the parameter for the αSyn group compared to AD without αSyn pathology, while a downward arrow indicates a decrease compared to the AD group. An arrow indicates an effect confirmed with statistical tests, while an arrow inside brackets suggests a potential effect not confirmed by statistical tests. ns – not significant

Other crucial findings of our study are the distinct neuropathological characteristics (Fig. 6C) observed between AD patients with two distinct distribution patterns of αSyn pathology. In our cohort, about half of the AD patients displayed the amygdala-predominant variant, while the other half exhibited either a caudo-rostral variant or a uniformly severe distribution of αSyn pathology across different brain regions. AD patients with the caudo-rostral αSyn pathology pattern, in contrast to those with the amygdala-predominant variant, are marked by lower severity of neuritic plaques and hippocampal pTau, and less prevalent CAA type 1. Other studies have suggested a relationship between CAA type 1, which is characterized by capillary involvement, and increased blood-brain barrier permeability.^71^ Moreover, it has been shown that severe CAA may contribute to the formation of microinfarcts^24^ and microhemorrhages^72^ and potentially contribute to neuronal degeneration. The question of whether the difference that we observe between the two αSyn pathology spreading patterns results from the different regions of pathology onset or from distinct biochemical properties, as some studies have suggested,^73^ requires further exploration.

Defining the distinct neuropathological characteristics of αSyn pathology spreading variants in AD patients could contribute to a better understanding of the LBD spectrum and improved patient stratification in the future. The recent advancements in α-Synuclein seeding assays^74–76^, such as the real-time quaking-induced conversion assay^77^, could play a pivotal role in patient tailored diagnostic approaches. While these assays have been extensively evaluated in cohorts with LBD, their diagnostic accuracy in AD patients remains unclear. Studies utilizing these assays have detected αSyn positivity in 30-45% of cases,^78,79^ slightly lower than the neuropathological estimation of 43-63%.^4–7^ This raises the question of whether these biomarkers enable the distinction between specific spreading patterns of αSyn pathology. For clinical purposes, differentiating between AD patients with various αSyn patterns is crucial, as αSyn interaction with other protein aggregates could impact clinical progression and treatment options for patients. Given the heterogeneity of pathological processes observed in AD patients with additional αSyn aggregates, treatment with anti-amyloid monoclonal antibodies^80^ may be less effective compared to other groups. Furthermore, the observation that AD patients with severe hippocampal degeneration often exhibit co-morbid αSyn and TDP-43 pathologies provides insights which could be utilized to improve diagnostic accuracy for patients in vivo, enhancing the reliability of clinical trials by better stratification.

This study has several limitations. Firstly, cases with an amygdala-predominant αSyn pathology pattern in our sample may be underdiagnosed, especially with low lesion burden. This is due to the highly heterogeneous nature of the amygdaloid complex and analyzing only a single representative section. Additionally, the dual-hypothesis of αSyn pathology spread assumes that the amygdala-predominant variant begins unilaterally,^81^ and our study only examined one hemisphere microscopically. Encouragingly, we observed a similar ratio of approximately 2-to-1 between cases with caudo-rostral and amygdala-predominant pattern, as Raunio,^13^ with a corresponding increase in amygdala-based αSyn pathology with increasing ADNC severity. This suggests that our method of stratification aligns with previous research. Another limitation is a result of strict exclusion criteria and stratifying a moderate-sized patient sample into three categories, which limited our statistical power to detect only large effects. We suspect that medium-sized effects, such as a higher pTau burden in AD with an amygdala-predominant αSyn pathology pattern compared to AD without αSyn lesions in the hippocampus, may have been overlooked. Similarly, our structural equation model satisfies the minimum criterion of 10 cases per estimate but falls short of the optimal standard of 20 cases per parameter estimate.^82^ Consequently, this limitation increases the likelihood of failing to detect minor effects.

In conclusion, our findings suggest that the presence of an amygdala-predominant αSyn pathology pattern in AD contributes indirectly to hippocampal neurodegeneration, potentially by enhancing the aggregation and toxicity of TDP-43 and pTau lesions. Our results emphasize the complexity of AD and supports the hypothesis that ‘co-pathologies’ such as TDP-43 or αSyn, are an integral part of AD in some patients and, by interacting with the hallmark pathologies, contribute to neuronal loss. The presence of two distribution patterns of αSyn pathology in AD, along with observed neuropathological differences in CA1 neuron loss and the presence of CAA type 1, highlights the necessity for precise patient stratification. This stratification should consider not only the molecular and morphological identity of co-pathologies but also the specific distribution pattern, which can result in local synergy effects or interactions with other protein aggregates.

## Supplementary Material

Supplementary files

## Acknowledgements

We express our gratitude to Bas Lahaje and Helena Ver Donck for their assistance with immunohistochemical staining. Special thanks to Sam Verrept for proposing the application of structural equation modeling to our neuropathological data. We also extend our appreciation to Grzegorz Walkiewicz, Bas Moonen, Jolien Schaeverbeke, and Annie Wiedmer for their insightful feedback on the project and help with drafting the manuscript.

