## Supplementary files for "Amygdala-predominant α-synuclein pathology exacerbates hippocampal neuron loss in Alzheimer’s disease"

Correspondence to:

### Supplementary Material

**Supplementary Figure 1** Representative photos of αSyn pathology burden for each semi-quantitative severity score

**Supplementary Figure 2** Scatterplots examining the linearity of the relationship between neuronal density in CA1 and numerical variables used in linear regression analysis

**Supplementary Figure 3** Illustration of the SEM employed in this study

**Supplementary Table 1** The clinical and neuropathological characteristics of the study cohort

**Supplementary Table 2** Exclusion criteria

**Supplementary Table 3** Antibodies used in this study

**Supplementary Table 4** Results of multiple linear regression models on CA1 neuronal density

**Supplementary Table 5** Parameters of multiple linear regression models on CA1 neuronal density

**Supplementary Table 6** Supplementary Table 6. Distribution of αSyn pathology severity scores across different brain areas in αSyn-positive cases

**Supplementary Table 7** Results from Dunn's Kruskal-Wallis multiple comparisons tests for αSyn-positive groups with varying ADNC

**Supplementary Table 8** Corrected P-values from Spearman's semi-partial correlation, with age at death as a covariate, for αSyn-positive cases

**Supplementary Table 9** Key differences between symptomatic AD patients without αSyn and with different spreading patterns of αSyn pathology

**Supplementary table 10** Results of ANOVA and Kruskal-Wallis tests on AD patients

**Supplementary table 11** Results of a post-hoc Dunn's multiple comparisons tests on AD patients

**Supplementary table 12** Results of pairwise-comparisons with Fisher's exact test on nominal variables on AD patients

**Supplementary Table 13** Results and fit statistics of the Structural Equation Model using observations from AD patients

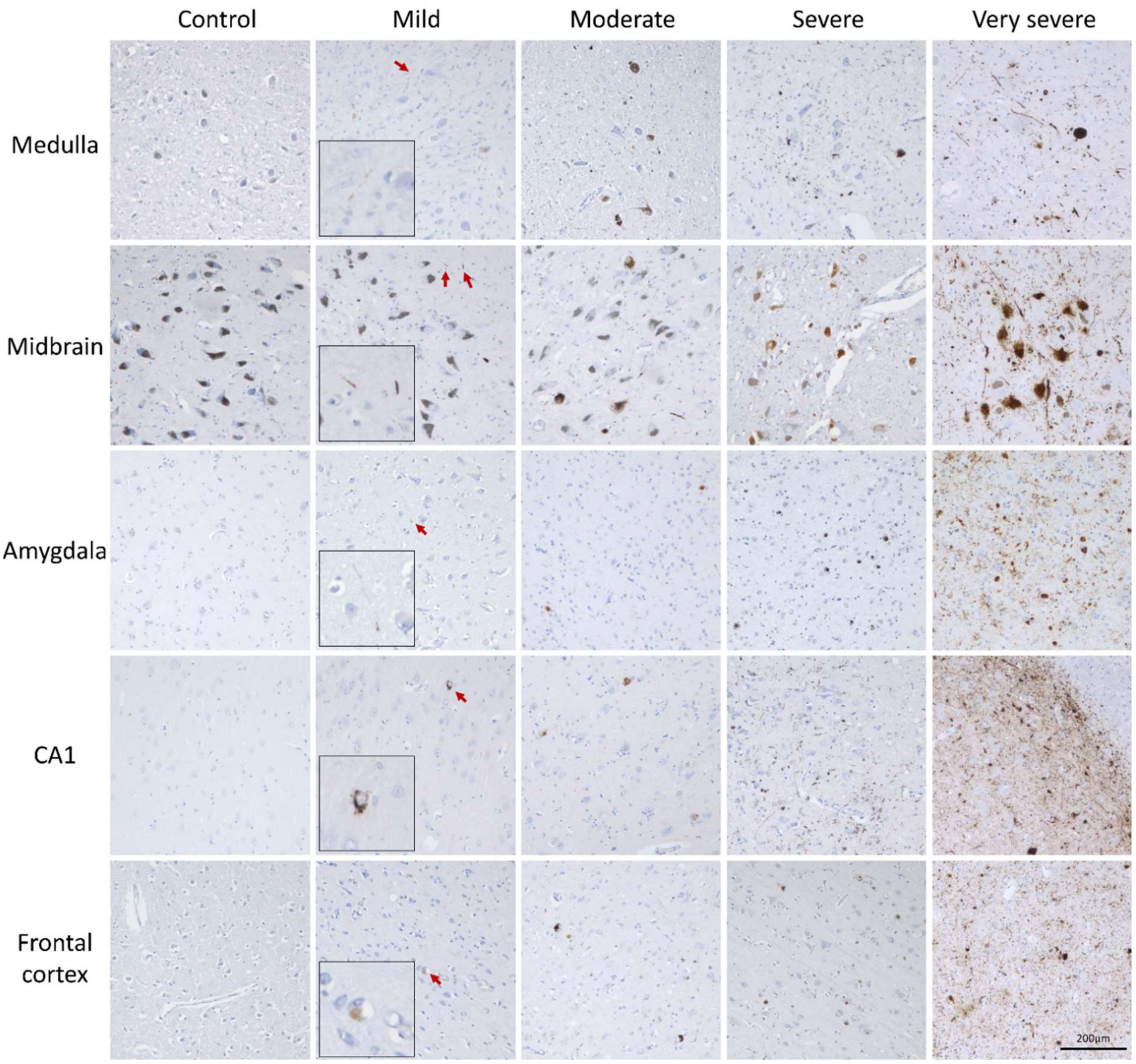

**Supplementary Figure 1 Representative photos of αSyn pathology burden for each semi-quantitative severity score** The graphic representation displays αSyn severity scores assigned to various brain regions, including the dorsal vagal nucleus in the medulla, substantia nigra in the midbrain, amygdaloid nucleus, CA1 subregion of the posterior hippocampus, and grey matter in the frontal cortex. Arrows in the mild condition indicate areas with αSyn pathology, which are further illustrated at higher magnification. The tissue was stained using the 5G4 antibody and photos were taken at a magnification of 200x.

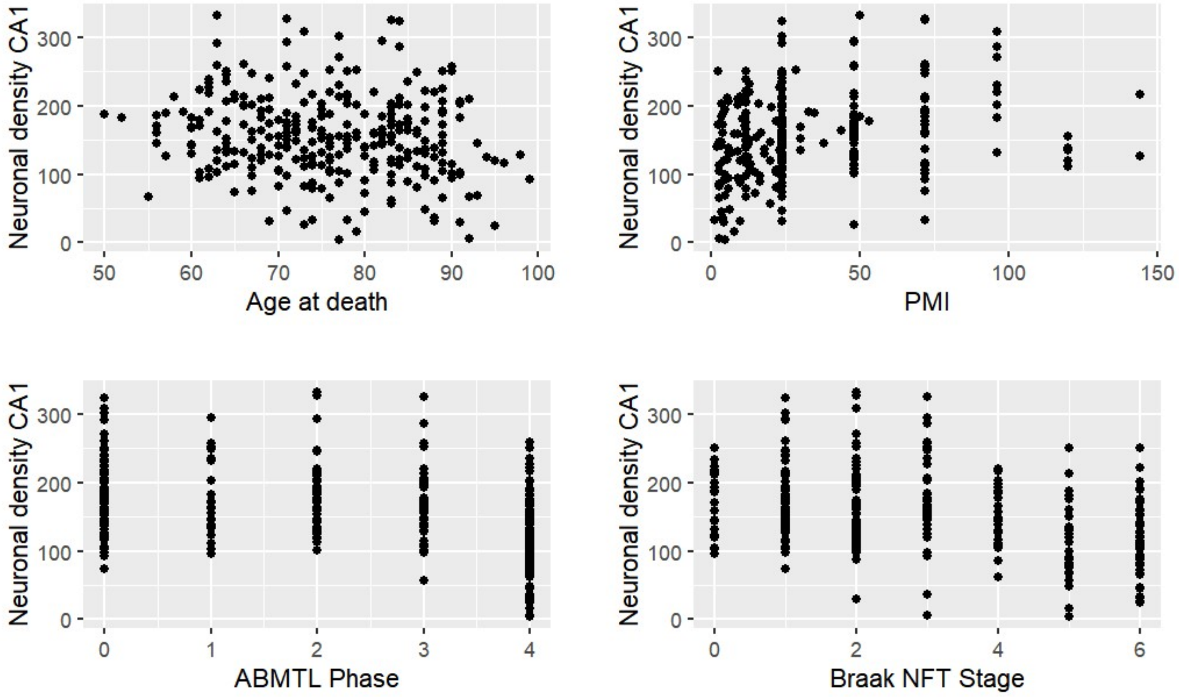

**Supplementary Figure 2 Scatterplots examining the linearity of the relationship between neuronal density in CA1 and numerical variables used in linear regression analysis** PMI – post mortem interval, ABMTL Phase – Aβ Phase in the MTL region

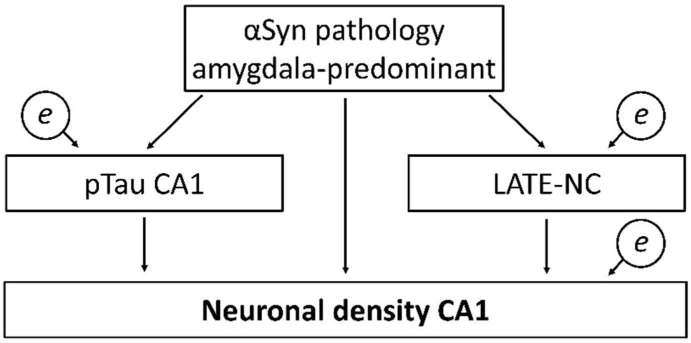

**Supplementary Figure 3 Illustration of the SEM employed in this study** The exogenous observed variable is amygdala-predominant αSyn pathology, while the endogenous observed variables include pTau in CA1, LATE-NC, and neuronal density in CA1. For each endogenous variable, the residual error term (depicted as small circles) has been estimated to account for unexplained variance. Arrows denote regression paths linking the variables, indicating the direction of influence from the independent to the dependent variable.

**Supplementary Table 1 The clinical and neuropathological characteristics of the study cohort (*n* = 291)**

| | | αSyn pathology negative | | | | αSyn pathology positive | | | |
|---|---|---|---|---|---|---|---|---|---|
| | | No ADNC (*n* = 45) | Mild ADNC (*n* = 69) | Moderate ADNC (*n* = 31) | Severe ADNC (*n* = 34) | No ADNC (*n* = 13) | Mild ADNC (*n* = 38) | Moderate ADNC (*n* = 23) | Severe ADNC (*n* = 38) |
| *Mean (SD)* | Age at death | 69.7 (9.7) | 73.8 (9) | 80.3 (10) | 79.5 (9.9) | 70.9 (7.2) | 76.7 (9.5) | 82.9 (8.2) | 77.1 (8.7) |
| | CDR | 0.1 (0.4) | 0.2 (0.6) | 1.6 (1.3) | 2.5 (0.7) | 0.3 (0.6) | 0.8 (1) | 2 (1.3) | 2.7 (0.6) |
| | Braak NFT Stages | 1.1 (1) | 1.7 (0.8) | 3.8 (0.7) | 5.7 (0.4) | 1.7 (1.2) | 1.8 (0.8) | 3.7 (0.8) | 5.6 (0.5) |
| | Aβ Phases MTL | 0 (0) | 1.8 (0.9) | 3.5 (0.7) | 3.9 (0.3) | 0 (0) | 2.1 (1.2) | 3.5 (0.6) | 4 (0.2) |
| | CERAD | 0 (0) | 0.1 (0.3) | 1.5 (0.9) | 2.8 (0.4) | 0 (0) | 0.4 (0.8) | 1.5 (0.7) | 2.5 (0.5) |
| | Neuronal density CA1 | 178.1 (57.4) | 182.6 (53.7) | 154.5 (61.2) | 127 (53.9) | 158.1 (29.8) | 160.8 (53.5) | 146.4 (62.6) | 104.5 (53.6) |
| | Braak PD Stages | 0 (0) | 0 (0) | 0 (0) | 0 (0) | 3.2 (1.8) | 4 (1.9) | 4.2 (1.8) | 4.4 (1.7) |
| % of cases | AP αSyn | 0% | 0% | 0% | 0% | 0% | 11% | 30% | 61% |
| | Sex (Male) | 60% | 48% | 42% | 44% | 77% | 66% | 43% | 58% |
| | Parkinsonism | 0% | 1% | 3% | 6% | 23% | 26% | 4% | 8% |
| | Epilepsy | 4% | 1% | 6% | 18% | 8% | 5% | 4% | 3% |
| | Dementia | 0% | 4% | 48% | 97% | 15% | 37% | 70% | 97% |

The mean and standard deviations (SD) of factors regarded as continuous variables are in the upper panel whereas nominal variables are summarized by % of cases with such characteristics. AP αSyn – amygdala-predominant αSyn pathology

**Supplementary Table 2 Exclusion criteria**

- Age of death <50y.o
- Presence of tumors or cancers in the central nervous system
- Central nervous system infections
- Severe brain edema or hypoxia
- Diagnosis of neurodegenerative conditions (Multiple Sclerosis, Huntington's Disease, FTLD, Multiple System Atrophy)

**Supplementary Table 3 Antibodies used in this study**

| | Antibody name | Source | Clonality; Host | Dilution | Stained regions |
|---|---|---|---|---|---|
| Aβ | Aβ17-24 (4G8) | BioLegend, US | Monoclonal; Mouse | 1:5000 | anterior MTL, occipital cortex |
| pTau | pTau Ser202, Thr205 (AT8) | Thermo Fisher, US | Monoclonal; Mouse | 1:1000 | anterior MTL, occipital cortex |
| αSyn | Anti-Aggregated αSyn (5g4) | Merck Millipore, Germany | Monoclonal; Mouse | 1:2000 | medulla, anterior MTL (if positive: brainstem, pons, posterior MTL, frontal cortex) |
| pTDP-43 | pTDP-43 409/410 | Cosmobio Co. LTD, Japan | Polyclonal, Rabbit | 1:5000 | anterior MTL, posterior MTL (if positive: frontal cortex) |

**Supplementary Table 4 Results of multiple linear regression models on CA1 neuronal density**

| | Model 1 | | | | Model 2 | | | | Model 3 | | | |
|---|---|---|---|---|---|---|---|---|---|---|---|---|
| | *β[CI 95%]* | *SE* | *t* | *P* | *β[CI95%]* | *SE* | *t* | *P* | *β[95%CI]* | *SE* | *t* | *P* |
| (Intercept) | 185.1 [132.4-237.9] | 26.8 | 6.9 | 3.25e-11 | 188.8 [136-241.5] | 26.8 | 7 | 1.51e-11 | 183.3 [125.1-241.5] | 29.6 | 6.2 | 2.26e-09 |
| Age at death | 0.1 [-0.6-0.8] | 0.4 | 0.3 | 0.749 | 0.0 [-0.7-0.7] | 0.4 | 0 | 0.980 | 0.0 [-0.7-0.8] | 0.4 | 0 | 0.985 |
| Sex (male) | 10.2 [-2.9-23.4] | 6.7 | 1.5 | 0.127 | 8.2 [-5-21.3] | 6.7 | 1.2 | 0.223 | 7.1 [-6.6-20.8] | 7 | 1 | 0.306 |
| Braak NFT stages | -7.1 [-12--2.1] | 2.5 | -2.8 | 0.005** | -6.3 [-11.4--1.2] | 2.6 | -2.4 | 0.016* | -3.8 [-9.2-1.6] | 2.7 | -1.4 | 0.167 |
| Aβ phases MTL | -6.9 [-12.8--1] | 3 | -2.3 | 0.022* | -5.3 [-11.3-0.6] | 3 | -1.8 | 0.079 | -6.5 [-12.7--0.3] | 3.1 | -2.1 | 0.041* |
| αSyn positivity | -18.0 [-31.6--4.4] | 6.9 | -2.6 | 0.010* | | | | | | | | |
| αSyn-AP | | | | | -39.6 [-61.5--17.8] | 11.1 | -3.6 | 4.11e-04*** | -42.8 [-65.5--20] | 11.6 | -3.7 | 2.62e-04*** |
| αSyn-CR | | | | | -5.4 [-20.6-9.7] | 7.7 | -0.7 | 0.483 | -7.2 [-22.7-8.3] | 7.9 | -0.9 | 0.360 |
| PMI | | | | | | | | | 0.2 [-0.1-0.5] | 0.1 | 1.5 | 0.142 |
| Hemisphere | | | | | | | | | -1.8 [-20.7-17] | 9.6 | -0.2 | 0.847 |
| Epilepsy | | | | | | | | | -34.0 [-63.2--4.8] | 14.8 | -2.3 | 0.023* |

αSyn AP – amygdala-predominant αSyn pathology variant, αSyn CR – caudo-rostral αSyn pathology variant, PMI – postmortem interval, hemisphere – hemisphere from which tissue has been taken; P-values have not been corrected for multiple testing. *<0.05, **<0.01, ***<0.001

**Supplementary Table 5 Parameters of multiple linear regression models on CA1 neuronal density**

| | **Model 1** | **Model 2** | **Model 3** |
|---|---|---|---|
| Sample size | 290 | 283 | 262 |
| Independent variables | age at death, sex, ABMTL Phase, Braak NFT Stage, αSyn+ | age at death, sex, ABMTL Phase, Braak NFT Stage, αSyn AP, αSyn CR | age at death, sex, ABMTL Phase, Braak NFT Stage, αSyn AP, αSyn CR, PMI, epilepsy, hemisphere |
| Model statistics | $F(5, 284) = 13.67$, $P = 5.82e\text{-}12$ | $F(6, 276) = 11.83$, $P = 8.32e\text{-}12$ | $F(9, 252) = 8.16$, $P = 1.25e\text{-}10$ |
| R2/Adjusted R2 | 0.194/0.18 | 0.205/0.187 | 0.226/0.198 |
| <u>Assumptions validation</u> | | | |
| Heteroscedasticity (Breusch-Pagan test) | $BP = 5.343$, $df = 5$, $P = 0.376$ | $BP = 4.962$, $df = 6$, $P = 0.549$ | $BP = 5.816$, $df = 9$, $P = 0.758$ |
| Normality of residuals (Shaphiro-Wilk test) | $P = 0.002$ | $P = 0.001$ | $P = 0.004$ |
| Variance Inflation Factor | all lower than 3 | all lower than 3 | all lower than 3 |

The models met all of the assumptions except for normality of residuals. We don't expect failing to meet his assumption to impact our results to great extent as our sample size could be considered more than 10 observations per variable[81]. αSyn AP – amygdala-predominant αSyn pathology variant, αSyn CR – caudo-rostral αSyn pathology variant

**Supplementary Table 6 Distribution of αSyn pathology severity scores across different brain areas in αSyn-positive cases (*n* = 105)**

| | Medulla | Midbrain | Amygdala | Temporal | CA1 | Frontal |
|---|---|---|---|---|---|---|
| No αSyn | 17 | 27 | 24 | 43 | 57 | 70 |
| Mild αSyn | 27 | 19 | 13 | 16 | 24 | 17 |
| Moderate αSyn | 21 | 24 | 21 | 17 | 9 | 11 |
| Severe αSyn | 23 | 18 | 18 | 17 | 9 | 4 |
| Very severe αSyn | 17 | 17 | 29 | 12 | 6 | 3 |

**Supplementary Table 7 Results from Dunn's Kruskal-Wallis multiple comparisons tests for αSyn-positive groups with varying ADNC (*n* = 105)**

| | αSyn Global Burden Score | | | αSyn MTL to brainstem ratio | | |
|---|---|---|---|---|---|---|
| Comparison (ADNC levels) | *Z* | *P.unadj* | *P.adj* | *Z* | *P.unadj* | *P.adj* |
| none - mild | -1.550 | 0.121 | 0.727 | -1.275 | 0.202 | 0.202 |
| none - moderate | -1.302 | 0.193 | 0.386 | -2.555 | 0.011 | 0.021* |
| mild - moderate | 0.205 | 0.838 | 1.000 | -1.743 | 0.081 | 0.098 |
| none - severe | -1.317 | 0.188 | 0.564 | -4.499 | <0.001 | <0.001*** |
| mild - severe | 0.338 | 0.736 | 1.000 | -4.349 | <0.001 | <0.001*** |
| moderate - severe | 0.097 | 0.923 | 0.923 | -2.133 | 0.033 | 0.049* |

*<0.05, **<0.01, ***<0.001

**Supplementary Table 8 Corrected P-values from Spearman's semi-partial correlation, with age at death as a covariate, for αSyn-positive cases (*n* = 105)**

| | Neuronal density CA1 | αSyn Medulla | αSyn Midbrain | αSyn Amy | αSyn Temp | αSyn CA1 | αSyn Frontal | αSyn GBS | αSyn AP |
|---|---|---|---|---|---|---|---|---|---|
| Neuronal density CA1 | 1 | 0.039 | 0.888 | 0.005 | 0.034 | 0.121 | 0.140 | 0.349 | 2E-05 |
| Medulla | 0.039 | 1 | 6.74E-14 | 0.002 | 9.39E-05 | 2.52E-06 | 6.01E-08 | 6.74E-14 | 9.27E-07 |
| Midbrain | 0.888 | 6.74E-14 | 1 | 1.69E-15 | 3.86E-15 | 3.37E-14 | 3.76E-15 | 3.86E-35 | 0.423 |
| Amygdala | 0.005 | 0.002 | 1.69E-15 | 1 | 1.84E-28 | 4.29E-19 | 7.34E-12 | 9.95E-31 | 4.16E-06 |
| Temporal | 0.034 | 9.39E-05 | 3.86E-15 | 1.84E-28 | 1 | 9.2E-30 | 1.91E-14 | 5.21E-35 | 2.34E-04 |
| CA1 | 0.121 | 2.52E-06 | 3.37E-14 | 4.29E-19 | 9.2E-30 | 1 | 2.57E-15 | 9.2E-30 | 0.024 |
| Frontal | 0.140 | 6.01E-08 | 3.76E-15 | 7.34E-12 | 1.91E-14 | 2.57E-15 | 1 | 1.03E-21 | 0.890 |
| αSyn GBS | 0.349 | 6.74E-14 | 3.86E-35 | 9.95E-31 | 5.21E-35 | 9.2E-30 | 1.03E-21 | 1 | 0.258 |
| αSyn AP | 2E-05 | 9.27E-07 | 0.423 | 4.16E-06 | 2.34E-04 | 0.024 | 0.890 | 0.258 | 1 |

AP – Amygdala-predominant αSyn pathology variant, αSyn GBS – αSyn pathology global burden score

**Supplementary Table 9 Key differences between symptomatic AD patients without αSyn and with different spreading patterns of αSyn pathology**

| | **no αSyn (*n* = 48)** | **Amygdala-predominant (*n* = 29)** | **Caudo-rostral (*n* = 22)** | **Observations per variable** |
|---|---|---|---|---|
| Age at death | 80.67 (9.62) | 78.21 (9.84) | 80.14 (7.09) | 99 |
| CDR | 2.46 (0.81) | 2.83 (0.58) | 2.72 (0.67) | 77 |
| αSyn global burden score | 0 (0) | 9.38 (5.4) | 9.91 (7.78) | 99 |
| Neuronal density CA1 | 126.88 (52.86) | 99.47 (52.04) | 129.84 (49.65) | 99 |
| Aβ Phases MTL | 3.77 (0.56) | 3.96 (0.19) | 3.68 (0.57) | 98 |
| CERAD | 2.56 (0.65) | 2.38 (0.73) | 2.14 (0.64) | 99 |
| Braak NFT Stages | 5.25 (0.89) | 5.24 (0.99) | 4.86 (1.04) | 99 |
| pTau CA1 | 0.51 (0.23) | 0.6 (0.24) | 0.39 (0.21) | 95 |
| LATE-NC | 0.7 (0.89) | 1.43 (1.07) | 1.05 (1.16) | 92 |
| CAA Severity | 1.73 (0.57) | 1.79 (0.49) | 1.59 (0.67) | 99 |
| CAA Type 1 | 65% | 69% | 32% | 99 |
| pTDP DG | 11% | 38% | 23% | 97 |
| Parkinsonism | 6% | 7% | 9% | 99 |
| Sex (Male) | 42% | 52% | 64% | 99 |

Variables considered as continuous have been summarized using the mean value and standard deviation (SD), while for nominal variables, we presented the percentage of cases with a positive outcome.

**Supplementary table 10 Results of ANOVA and Kruskal-Wallis tests on AD patients**

| Test | Variable | Summary |
|---|---|---|
| ANOVA | pTau CA1 | $F(2,92) = 5.27$, $P = 0.007$ |
| ANOVA | Neuronal density CA1 | $F(2,96) = 3.08$, $P = 0.051$ |
| ANOVA | Age at death | $F(2,96) = 0.66$, $P = 0.518$ |
| kruskal.test | αSyn global burden score | $H(2) = 82.97$, $P = 9.64e-19$ |
| kruskal.test | LATE-NC | $H(2) = 8.1$, $P = 0.0174$ |
| kruskal.test | CERAD | $H(2) = 7.14$, $P = 0.028$ |
| kruskal.test | CDR | $H(2) = 5.38$, $P = 0.068$ |
| kruskal.test | Aβ Phases MTL | $H(2) = 5.33$, $P = 0.070$ |
| kruskal.test | Braak NFT Stages | $H(2) = 2.63$, $P = 0.269$ |
| kruskal.test | CAA Severity | $H(2) = 2.1$, $P = 0.351$ |

Statistical test was applied to detect differences in parameters among AD patients without αSyn pathology and with two αSyn pathology spreading patterns

**Supplementary table 11 Results of a post-hoc Dunn's multiple comparisons tests for AD patients**

| Variable | Comparison | *Z* | *P.unadj* | *P.adj* | |
|---|---|---|---|---|---|
| Age at death | AP x αSyn- | -1.05 | 0.292 | 0.875 | |
| Age at death | AP x CR | -0.48 | 0.633 | 0.949 | |
| Age at death | αSyn- x CR | 0.44 | 0.662 | 0.662 | |
| αSyn global burden score | AP x αSyn- | 7.79 | 6.80e-15 | 2.04e-14 | *** |
| αSyn global burden score | AP x CR | 0.00 | 1.000 | 1.000 | |
| αSyn global burden score | αSyn- x CR | -7.11 | 1.12e-12 | 1.68e-12 | *** |
| Aβ Phases MTL | AP x αSyn- | 1.57 | 0.117 | 0.176 | |
| Aβ Phases MTL | AP x CR | 2.27 | 0.023 | 0.070 | . |
| Aβ Phases MTL | αSyn- x CR | 1.06 | 0.289 | 0.289 | |
| Braak NFT Stages | AP x αSyn- | 0.09 | 0.931 | 0.931 | |
| Braak NFT Stages | AP x CR | 1.43 | 0.153 | 0.230 | |
| Braak NFT Stages | αSyn- x CR | 1.49 | 0.136 | 0.409 | |
| CAA Severity | AP x αSyn- | 0.71 | 0.480 | 0.480 | |
| CAA Severity | AP x CR | 1.45 | 0.148 | 0.444 | |
| CAA Severity | αSyn- x CR | 0.94 | 0.345 | 0.518 | |
| CDR | AP x αSyn- | 2.22 | 0.027 | 0.080 | . |
| CDR | AP x CR | 0.56 | 0.574 | 0.574 | |
| CDR | αSyn- x CR | -1.44 | 0.151 | 0.226 | |
| CERAD | AP x αSyn- | -1.14 | 0.252 | 0.252 | |
| CERAD | AP x CR | 1.47 | 0.141 | 0.212 | |
| CERAD | αSyn- x CR | 2.66 | 0.008 | 0.023 | * |
| LATE-NC | AP x αSyn- | 2.84 | 0.005 | 0.013 | * |
| LATE-NC | AP x CR | 1.35 | 0.177 | 0.266 | |
| LATE-NC | αSyn- x CR | -1.13 | 0.258 | 0.258 | |
| Neuronal density CA1 | AP x αSyn- | -2.20 | 0.028 | 0.084 | . |
| Neuronal density CA1 | AP x CR | -1.87 | 0.061 | 0.092 | . |
| Neuronal density CA1 | αSyn- x CR | -0.05 | 0.959 | 0.959 | |
| pTau CA1 | AP x αSyn- | 1.35 | 0.178 | 0.178 | |
| pTau CA1 | AP x CR | 3.16 | 0.002 | 0.005 | ** |
| pTau CA1 | αSyn- x CR | 2.24 | 0.025 | 0.038 | * |

Tests were performed for variables that reached significance with ANOVA or Kruskal Wallis tests; αSyn- – AD αSyn-negative, AP – AD with amygdala-predominant αSyn pathology, CR – AD with caudo-rostral αSyn pathology; P-values have been adjusted with the Benjamini–Hochberg method per variable;
. <0.1, *<0.05, **<0.01, ***<0.001

**Supplementary table 12 Results of pairwise-comparisons with Fisher's exact test on nominal variables on AD patients**

| Variable | Comparision | *P.unadj* | *P.adj* | |
|---|---|---|---|---|
| CAA Type I | AP x CR | 0.012 | 0.028 | * |
| CAA Type I | αSyn- x AP | 0.805 | 0.805 | |
| CAA Type I | αSyn- x CR | 0.019 | 0.028 | * |
| Parkinsonism | AP x CR | 1.000 | 1.000 | |
| Parkinsonism | αSyn- x AP | 1.000 | 1.000 | |
| Parkinsonism | αSyn- x CR | 0.646 | 1.000 | |
| pTDP DG | AP x CR | 0.362 | 0.362 | |
| pTDP DG | αSyn- x AP | 0.009 | 0.026 | * |
| pTDP DG | αSyn- x CR | 0.316 | 0.362 | |
| Sex (Male) | AP x CR | 0.569 | 0.569 | |
| Sex (Male) | αSyn- x AP | 0.480 | 0.569 | |
| Sex (Male) | αSyn- x CR | 0.123 | 0.369 | |

αSyn- – AD αSyn-negative, AP – AD with amygdala-predominant αSyn pathology, CR – AD with caudo-rostral αSyn pathology; P-values have been adjusted with the Benjamini–Hochberg method per variable;
. <0.1, *<0.05, **<0.01, ***<0.001

**Supplementary Table 13 Results and fit statistics of the Structural Equation Model using observations from AD patients (*n* = 90)**

| | | | | | |
|---|---|---|---|---|---|
| **Regressions** | | | | | |
| | *Estimate* | *Std. Estimate* | *SE* | *z* | *P* |
| LATE-NC ~ | | | | | |
| αSyn AP | 0.622 | 0.276 | 0.228 | 2.724 | 0.006 |
| pTau CA1 ~ | | | | | |
| αSyn AP | 0.11 | 0.221 | 0.051 | 2.15 | 0.032 |
| Neuronal Density CA1 ~ | | | | | |
| pTau CA1 | -81.139 | -0.357 | 19.9 | -4.077 | <0.001 |
| LATE-NC | -21.261 | -0.422 | 4.472 | -4.754 | <0.001 |
| αSyn AP | -5.318 | -0.047 | 10.317 | -0.515 | 0.606 |
| **Variances** | | | | | |
| | *Estimate* | *Std. Estimate* | *SE* | *z* | *P* |
| LATE-NC | 1.006 | 0.924 | 0.15 | 6.708 | < 0.001 |
| pTau CA1 | 0.051 | 0.951 | 0.008 | 6.708 | < 0.001 |
| Neuronal Density CA1 | 1810.945 | 0.656 | 269.96 | 6.708 | < 0.001 |
| **Fit statistics** | | | | | |
| Measure | Obtained values | Good fit criterium[60] | | | |
| Test statistic | 0.023 | | | | |
| Degrees of freedom | 1 | | | | |
| *P* (Chi-square) | 0.88 | >0.05 | | | |
| Comparative Fit Index (CFI) | 1 | >0.95 | | | |
| Tucker-Lewis Index (TLI) | 1 | >0.95 | | | |
| RMSEA [95% CI] | 0 [0, 0.14] | [0, 0.08] | | | |
| SRMR | 0.006 | <0.05 | | | |

AP – amygdala-predominant variant of αSyn pathology, SE – standard error, Std – standardized values
